# Dark adaptation is not necessary to drive the proliferative regeneration response in the adult zebrafish retina

**DOI:** 10.1101/2023.08.28.554932

**Authors:** Marlo Hemerson, Jeremy Muhr, Lauren Pferdmenges, Sarah Jiudice, Kristin M. Ackerman

**Affiliations:** Department of Biology, Program in Neuroscience, High Point University, High Point, North Carolina, USA; Department of Biology, High Point University, High Point, North Carolina, USA; Department of Medical Sciences, High Point University, High Point, North Carolina, USA

## Abstract

Macular degeneration, retinal detachment, and retinitis pigmentosa are eye diseases which cause damage to the photoreceptors and are considered leading causes of vision loss worldwide (National Eye Institute, 2022). Thus, developing animal models to study processes related to retinal damage, specifically loss of photoreceptors and their pending degeneration/regeneration is critical. This study aims to modify current light-induced retinal degeneration (LIRD) protocols within the zebrafish model to more efficiently replicate the proliferative response that drives regeneration. The most common LIRD model incorporates a 10-14 day dark adaptation period followed by multiple days of constant intense light exposure designed to damage the retinas and induce a proliferation response (Thomas et al., 2012; Vihtelic & Hyde, 2000). Modified procedures have been published and demonstrate that shortened 24-hour dark adaptation periods still result in cellular damage via apoptosis, but the proliferation response (i.e., pending regeneration) was not examined (Khan et al., 2020). Therefore, our study aims to fully eliminate dark adaptation and will use changes in retinal morphology and proliferation as the hallmark of a successfully initiated regeneration response. Dark-adapted (DA-control fish) and non-dark-adapted (NDA-experimental fish) were taken off their housing system and exposed to constant intense light for 24, 36, and 48 hr. to ensure enough time for damage and proliferation to proceed. Retinas were cross-sectioned to include dorsal and ventral retina, immunohistochemically labeled, and the outer nuclear layer (ONL), inner nuclear layer (INL), cones, and proliferation cell nuclear antigen (PCNA)-positive cells were analyzed to determine the level of structural integrity, if cell photoreceptor numbers were altered, and if proliferation occurred in response to light exposure. Dark-adapted (DA) and non-dark-adapted (NDA) zebrafish retinas displayed similar damage and increased proliferation responses to the constant intense light, indicating that the 10–14-day dark adaptation period is not necessary to induce regeneration in adult zebrafish retinas, reducing experimental timing by two weeks.

## Introduction

Degenerative retinal diseases, including but not limited to macular degeneration, retinal pigmentosa, and diabetic retinopathy are characterized by damage to photoreceptors or ganglion cells within the retina, a tissue layer lining the back of the eye that is responsible for visual reception and transmission to the brain. Destruction of photoreceptor is the first step in the degenerative process, and it leads to degeneration of the rest of the retina, eventually causing blindness or severely limited sight (Lee et al., 2021). Due to the inability of human retinal tissue to regenerate, there is currently no known cure or treatment therapy capable of reversing this damage. Investigating potential mechanisms by which photoreceptor regeneration occurs in other animal models may provide insight into the development of retinal degeneration therapies.

Zebrafish are a suitable animal model for studying human optic disease because of their fully mapped genome, structurally similar optic features, and unique ability to regenerate retinal tissue (Chhetri et al., 2014; Noel et al., 2021). Zebrafish retinal regeneration is driven by Müller glia, or retinal stem cells, which can generate multipotent retinal progenitor cells to replace lost/damage cells (Gemberling et al., 2013). Current research suggests that the mechanism by which Müller glia can re-enter the cell cycle and differentiate into photoreceptors is due to the regulation of transcription factors, genetic expression, and microenvironmental niches (Raymond et al., 2006). In response to cell death and inflammation, cells release various growth factors that initiate Müller glia proliferation, and through signaling cascades, result in Müller glia dedifferentiation. Dedifferentiation is quickly followed by the proliferation of progenitor cells that can differentiate and develop into mature photoreceptors (Raymond et al., 2006; Thummel et al., 2009). In the uninjured retina, glia migrate to the INL and divide symmetrically, thus continuing self-renewal of the stem cell population. Following photoreceptor lesion (light damage), however, injured Müller glia are reprogrammed to divide asymmetrically upon migration to the INL. Contrarily, this asymmetric division produces multipotent retinal progenitor cells (Lahne & Hyde, 2017).

Retinal regeneration is marked by Müller glia re-entering the cell cycle and commonly visualized by the expression or upregulation of proliferating cell nuclear antigen (PCNA) (Thummel et al., 2009). Current understanding of the mechanisms by which this retinal regeneration occurs is largely due to an experimentation protocol involving a period of dark adaption and a light-damage lesion, followed by immunohistochemistry (IHC) staining for PCNA at various time points to observe upregulation of progenitor cell proliferation. Traditional light-induced retinal degeneration (LIRD) paradigms begins (Figure 1 D) with 14 hr. light:10 hr. dark standard light care cycle (SLC-Figure 1A), followed by the traditional experimentation protocol of 10-14 days of dark adaptation (DA-Figure 1 B), and finally constant intense light treatment (LT-Figure 1C) with immunohistochemistry data collection at various timepoints between 1-4 days depending on the experimental question (Thomas et al., 2012; Vihtelic & Hyde, 2000). While this method produces reliable, satisfactory results, the time needed to conduct one trial is lengthy. Current literature shows that this lengthy dark adaptation period of 14 days is not needed, and that 24 hr. of dark adaptation damages the retinas and induces cell death by apoptosis to the same extent as the 14-day period with photoreceptor regeneration by day 28 post-LIRD, but progenitor cell proliferation was not verified (Khan et al., 2020). Our study proposes an alternate method of experimentation in which SLC is directly followed by light damage (full elimination the dark adaptation period-Figure 1 E). Also, an attempt to understand the mechanism by which the regeneration occurs is presented through careful analysis of the proliferating PCNA-positive cells immediately following LIRD at 0, 24, 36, and 48 hr. periods in both dark-adapted (DA) and non-dark-adapted (NDA) zebrafish groups. We show here that dark adaption prior to light lesion does not appear to be a necessary component to structurally damage the photoreceptors layer and that the number of PCNA-positive cells following both the traditional (DA) and altered trial approach (NDA) increase similarly a successfully initiated regeneration response. These data contribute to a growing understanding of the experimental conditions necessary to induce regeneration and introduce a more streamlined, efficient method by which results can be obtained two weeks quicker.

**Figure 1:**
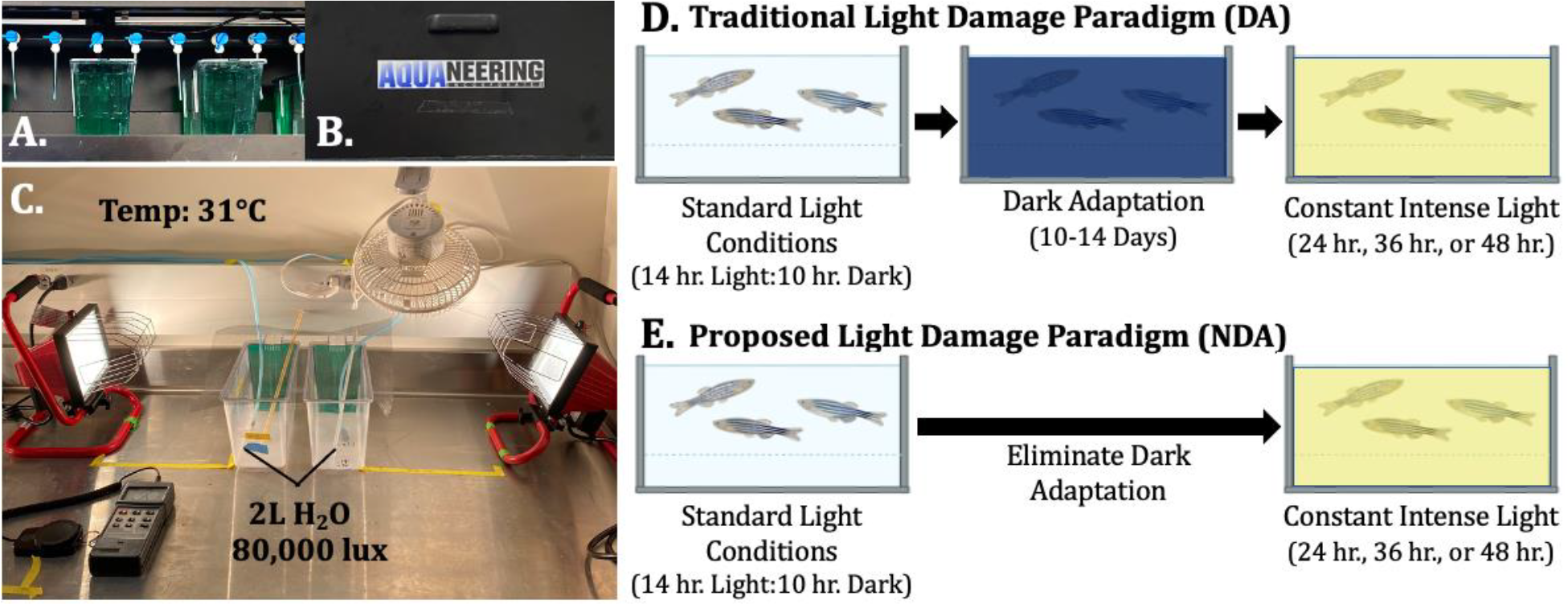
New Protocol for Light Damage Protocol with lacks Dark Adaptation. A,C,D) In traditional Light Damage Paradigms (DA), adult zebrafish are removed from Standard Light Conditions (A,D) are placed in a Light/Dark box (B-Aquaneering) for 10 to 14 days of Dark Adaptation (D) and then placed in Constant Intense Light (C,D) for damage. Our newly design protocol (NDA), removed adult zebrafish from Standard Light Conditions (A,E) and then placed directly in Constant Intense Light (C,E).

## Methods

### Animals

Adult male and female zebrafish (*Danio rerio*) were used for the study. Zebrafish were maintained under standard light conditions (14 hr. light:10 hr. dark cycle) at 29ºC. The experimentation followed High Point University guidelines approved by the Institutional Animal Care and Use Committee.

### Light-induced retinal degeneration model

For the dark-adapted protocols (DA), zebrafish were taken from the standard light conditions and placed in the light-tight box (Aquanering INC, cat. ZLB1060) for dark adaptation for 14 days (Figure 1A and 1B), while the non-dark adapted (NDA) zebrafish were kept under standard light conditions during this time (Figure 1A). After the 14 days (about 2 weeks), both DA and NDA zebrafish were placed into tanks with 2 L of system water surrounded by two portable halogen work lights (Walmart, SKU: ZX9JNSN59462 500 W T-3 Type Halogen Bulb) placed 10 in. from the tank (Figure 1C). The light intensity in the tank created by this setup was 80,000 lux and was measured using a Dual-Display Traceable Light Meter (Traceable Calibration Control Company, cat. 3252CC). Fish (n ≥ 10) were collected after 0, 24, 36, and 48 hr. of constant intense light exposure.

### Tissue preparation and cryosectioning

Zebrafish were euthanized with an overdose of 1:500 2-phenoxyethanol and enucleated eyes were fixed in 9:1 ethanolic formaldehyde overnight at 4°C. The eyes were then serially washed with ethanol (100%, 95%, 80%, 70%, 50%) for 5 min each at room temperature, then the retinas were transferred into serial sucrose washes (5%, 3 X 10 min and 30%, 1 X 4 hr.) at room temperature. A 2:1 tissue freezing medium (2 parts OTC to 1 part 30% sucrose solution) was added and the retinas remained in that overnight at 4°C. The next day, the retinas were placed in tissue freezing medium and frozen overnight at - 80°C to prepare for sectioning. Frozen eyes were vertically sectioned at the central portion of the retina using the cryostat (Leica, CM1860 UV) at 14um thickness and placed onto super frost charged slides, heated at 55°C for 4 hr., then stored at -80°C.

### Antibodies

Primary antibodies used in this study were mouse anti-ZPR-1 (1:250, Zebrafish International Research Center, ZDB-ATB-081002-43) to stain cone cells and rabbit anti-PCNA (1:1000, Sigma 8825) to stain proliferative cells. The secondary antibodies used for fluorescent detection were anti-mouse conjugated with Alexa Fluor 594 (1:500, Life Technologies, A11032) and anti-rabbit conjugated with Alexa Fluor 488 (1:500, Life Technologies, A11034). A nuclear fluorescent stain, 4’,6-Diamidino-2-Phenylindole, Dihydrochloride (DAPI) (1:10,000, Life Technologies, D1306) was used to stain cell nuclei for rod quantification.

### Immunohistochemistry

Frozen retinal sections were labeled using immunohistochemistry. The slides were washed three times with 1X phosphate buffered saline (PBS) (Sigma, P-5368, lot 049K8204) for 5 min. and then 1 hr. of blocking with a solution of 2% normal goat serum, 0.2% Triton X-100, 1% DMSO, and 1X PBS. Primary antibodies were added, and the slides were left overnight at room temperature to incubate in a humidity chamber. The following day, the slides were washed in 1X PBS 0.05% Tween-20 (PBS-T) solution three times for 10 min each (30 min. Total). The secondary antibodies were added at 1:500 and incubated at room temperature for 1 hr. in a dark humidity chamber. After, the slides underwent three additional washes for 10 min. with PBS-T. The second wash had the DAPI stain added to solution. A fourth and final wash with 1X PBS for 10 min. was performed. Prolong Gold was used to mount coverslips on slides. The finished slides remained overnight at room temperature to harden and were then stored at 4°C indefinitely to be quantified.

### Fluorescent imaging and analysis

Images were taken using an Invitrogen EVOS (Thermo-Fisher Scientific, M5000) microscope with a 20X objective. All cell quantification was taken over a 200-um area of the dorsal central retina. To measure outer nuclear layer (ONL) thickness, software analysis was utilized to draw a scale-bared line from the basal-most rod nuclei to the apical end of the cone nuclei. Within the 200-um area in the dorsal central retina, three ROI (one to the right, one to the left, and one in the middle of the 200-um area) of the ONL were averaged. Within the 200-um dorsal central area, the number of DAPI-labeled rod nuclei were counted by taking three 20-um sections (one to the right, one to the left, and one in the middle), counting each cell nuclei in the 20-um area, then the three sections were averaged. The total number of ZPR-1-positive cones and the PCNA-positive progenitor and precursor cells were counted across the entire 200 um area (ROI were not selected). The differences between groups were analyzed using one-way ANOVA with a Tukey post hoc test. Variance was reported as mean +/-SEM and significance was identified at p≤0.05. Celleste 5.0 Image Analysis Software (SKU# AMEP4865) was used to brighten images.

## Results

### Light-induced retinal degeneration significantly decreased ONL thickness and number of rod nuclei with and without dark adaptation

The traditional LIRD models (DA) causes structural damage, specifically causing decreased ONL thickness and increased pyknotic nuclei resulting from cell death (Raymond et al., 2006; Thomas et al., 2012; Thummel et al., 2009; Vihtelic & Hyde, 2000). To assess retinal morphology in our DA and NDA groups, retinas were labeled with DAPI to visual the ONL. Damage was defined as loss of ONL structure/integrity (Figure 2A), a thinning of the ONL (Figure 2B), and decreased number of DAPI-positive nuclei in the ONL (Figure 2C). Qualitatively, our experimental group of non-dark adaptation (NDA) displayed damage to the retinas similarly to the DA controls. The disorganized morphology and loss of ONL integrity is evident in both paradigms as early as 24 hours of constant intense light and continues through 48 hours (about 2 days) of damage. As seen in Figure 2A, the more time spent in LT, the less structured the ONL appeared. The retinas become thinner and more dispersed, and the cells themselves take on a shattered-looking effect resultant from the light damage, as compared to the tightly packed and organized 0 hr. retinas. We note that NDA rod nuclei had a unique shape compared to the DA rod nuclei, appearing as concaved with a thick rim and a light center. This shape first shows up at the 24-hr. time point and continues through 48 hr.

**Figure 2:**
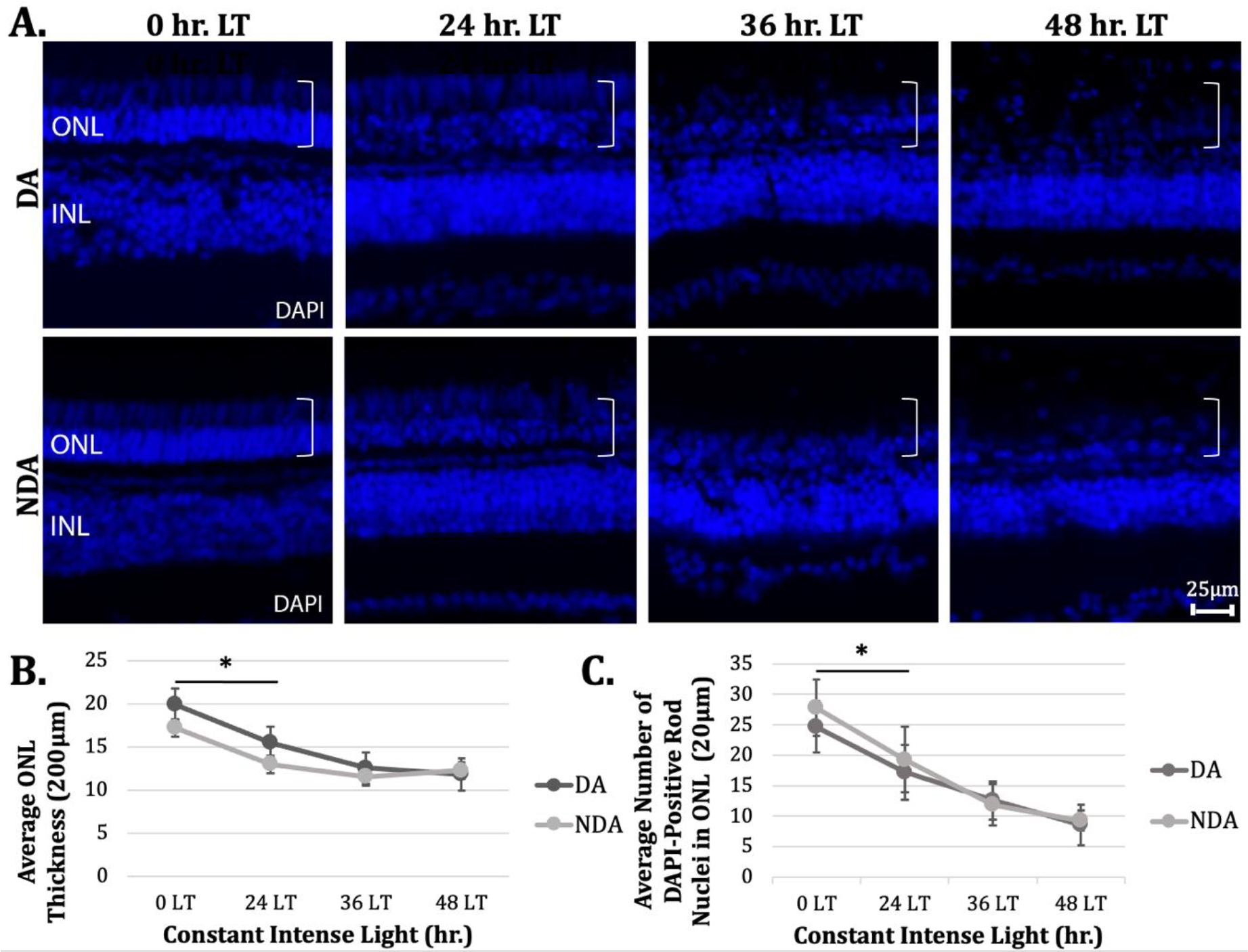
Light damage significantly decreased ONL thickness and number of rod nuclei in both dark-adapted and non-dark-adapted retinas. A) DA and NDA retinas were light damaged, sectioned, and counter stained with the nuclear dye, DAPI (blue). Light-damaged retinas exhibit a disorganization and collapse in the ONL as compared to the tightly packed/organized undamaged controls (white bar). The DA retinas are a positive control and demonstrate ONL damage beginning at 24 LT and complete by 36 hours LT. Disruption in the NDA retina is evident as early as 24 hours of LT. B) As an indicator of damage, quantification of ONL thickness was measured across 200 um; both DA and NDA exhibit a significant decrease in thickness as the retina damages. Importantly, there is no significant difference between DA and NDA indicating that both paradigms damage equally well. C) The decrease in ONL thickness was not only due to a change in cell morphology, but also a significant reduction in the number of DAPI-positive nuclei in both the DA and NDA conditions. Again, there was no significant difference between the DA and NDA conditions at any damage timepoint (0, 24, 36, or 48). n≥10, p≤.05, ANOVA, Tukey post hoc.

Although qualitative visualization of the retina demonstrates damage, we quantified ONL thickness changes (Figure 2B). The white line on the first photo block indicates an example measurement taken to evaluate the thickness of the ONL, measuring from the most basal rod photoreceptor nuclei to the apical edge of the cone nuclei. Comparing the DA to NDA retinas, both models had a significant decrease in the ONL thickness from 0 to 24 hr., indicating similar damage experienced in both paradigms. The DA retinas decreased from 19.9±4.2um at 0 hr. to 15.5±3.7um at 24 hr. (p=0.024), while the NDA retinas decreased from 17.2±4.3um at 0 hr. to 13.0±1.6um at 24 hr. (p=0.034). However, there was no significant difference between the groups at any timepoint (0 hr. [19.9±4.2um DA:17.2±4.3um NDA, p=0.40]; 24 hr. [15.5±3.7um DA:13.0±1.6um NDA, p=0.60]; 36 hr. [12.6±2.0um DA:11.5±2.3um NDA, p=0.96]; 48 hr. [11.8±2.2um DA:12.2±3.4um NDA, p=1.0]) indicating no difference in the models. Reduction in ONL thickness could be due to reduction in nuclei size or a reduction in photoreceptor numbers (rods or cones). First, the number of rod nuclei was across three 20-um sections to determine if the density of cells within the ONL changed. The number of rod nuclei significantly decreased across the time points in both paradigms, indicating no difference in dark adaptation for reducing the number of photoreceptors via damage. In the DA retinas, rod nuclei decreased from 24.7±4.2 cells to 17.2±4.5 cells (p=0.024) and the NDA retinal cells decreased from 27.8±4.6 cells to 19.3±5.4 cells (p=0.034) during the 0 hr. to 24 hr. time points. Again, there is no significant difference between DA and NDA groups at any time point (0 hr. [24.7±4.2 cells DA:27.8±4.6 cells NDA, p=0.47]; 24 hr. [17.2±4.5 cells DA:19.3±5.4 cells NDA, p=0.96]; 36 hr. [12.6±3.1 cells DA:11.9±3.4 cells NDA, p=1.0]; 48 hr. [8.5±3.3 cells DA:9.3±1.6 cells NDA, p=1.0]).

### Light-induced retinal degeneration significantly decreased cone cells with and without dark adaptation

It is evident from the previous figure that rod photoreceptors are decreasing in both DA and NDA paradigms, so our next objective was to examine cone photoreceptor health. Current literature regarding retinal cone cell response to light damage is inconclusive some studies showing a decrease and some studies showing no change (D’Orazi et al., 2020; Khan et al., 2020; Vihtelic & Hyde, 2000). To visualize the effect DA and NDA have on cone cells, the retinas were stained with ZPR-1, an antibody used to label red/green double cones. Qualitatively, figure 3A shows the unhealthy morphological progression as light damage increases; the 0 hr. cones of both DA and NDA groups look healthy, as demonstrated by their alignment, structured process extending apically and basally with end-feet visualized. As damage increases at 36 and 48 hr. LT, the cones fray at the ends and the end-feet disappear. To determine if there was a loss of cone numbers, total ZPR-1-positive cells were quantified across a 200-um area of the dorsal central retina. Like rod photoreceptors, the number of cones also significantly decreased from 0 to 24 hr. in both groups (DA [45.4±8.4 cells 0 hr:33.3±4.2 cells 24 hr, p=0.033]; NDA [38.5±10.2 cells 0 hr:18.4±7.9 cells 24 hr, p=0.1.12e-8]), but then plateaued to 32 cells in DA and 20 cells in NDA for the remaining time points. Between the two groups, there is no significant difference at 0 hr. or 48 hr. (0 hr. [45.4±8.4 cells DA:38.5±10.2 cells NDA, p=0.43]; 48 hr. [31.9±3.9 cells DA:22.4±2.9 cells NDA, p=0.12]); however, at the 24 and 36 hr. time points, there is a significant difference between DA and NDA (24 hr. [33.3±4.1 cells DA:18.4±7.9 cells NDA, p=0.00188]; 36 hr. [33.2±2.5 cells DA:19.4±9.7 cells NDA, p=0.00163]). Even with this significant difference, there is a similar pattern between the groups with a robust decline between 0 to 24 hr., then no change thereafter.

**Figure 3:**
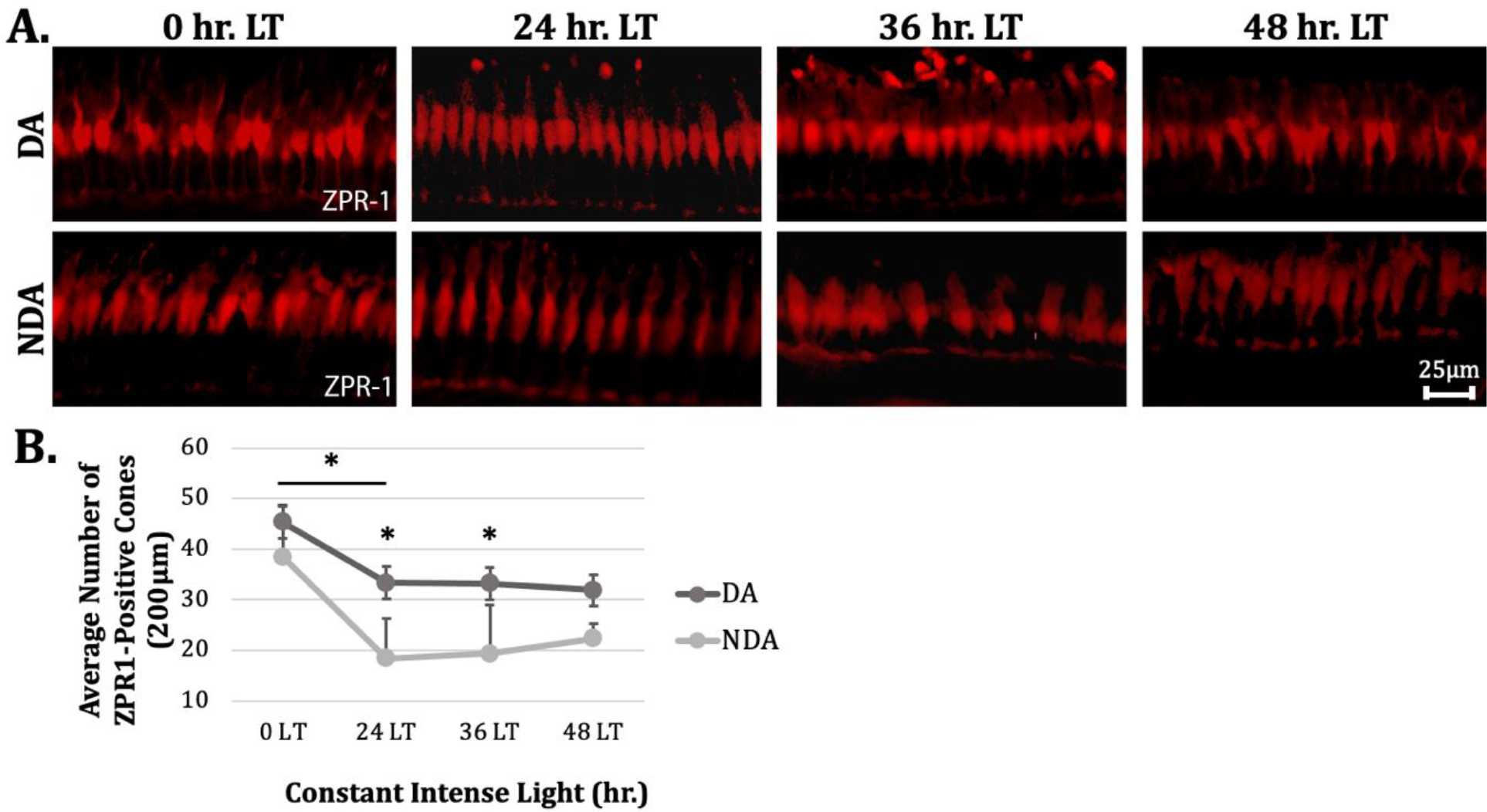
Light damage significantly decreased cone cells in both dark-adapted and non-dark-adapted retinas. A) DA and NDA retinas were light damaged, sectioned, and immunohistochemically labeled for ZPR-1 (red), an antibody that labels red/green double cones. B) Quantification demonstrates a significant loss in a subset of cone photoreceptors in light damaged retinas in both DA and NDA treatments as compared to undamaged controls. Interestingly, there is significant difference in the number of cones between the DA and NDA at 24 hr. and 36 hr. light treatment, with p=0.00188 and p=0.00163, respectively. n≥10, p≤.05, ANOVA, Tukey post hoc.

### Light-induced retinal regeneration significantly increased INL PCNA cells in both dark and non-dark adapted retinas

With reductions in both rod and cone photoreceptors, we can hypothesize that a proliferative regeneration response will initiate if enough damage was induced. The hallmark of retinal regeneration is the dedifferentiation of Müller glia to multipotent progenitor cells, followed by their asymmetrical division and migration from the INL to ONL to replace damaged cells. This regeneration-generated progenitor cell population can be labeled using antibodies against proliferation cellular nuclear antigen (PCNA) (Figure 4A). Literature indicates that LIRD will induce increased numbers of PCNA-positive cells in the INL as the progenitors are dividing before migrating to the originally damaged/lost photoreceptors. In our study, both the DA and NDA groups both significantly upregulated the number of PCNA-positive cells at 36 hr. LT compared to controls, indicating a successful regeneration response. However, there are discrepancies between DA and the new NDA paradigm (Figure 3B). NDA retinas have a much earlier and more robust proliferation response, with cell numbers significantly increasing from 0 to 24 hr. time points, while DA retinas show no increase in INL during this same period (DA [0 cells 0 hr:0.6±1.3 cells 24 hr, p=1.0]; NDA [0.19±0.5 cells 0 hr:12.1±9.1 cells 24 hr], p=0.00466]). The trend switches from the 24 to 36 hr. period, with DA retinas significantly increasing and NDA retinal proliferation staying stable (DA [0.6±1.3 cells 24 hr:14.6±5.9 cells 36 hr, p=6.27e-5]; NDA [12.1±9.1 cells 24 hr:13.8±7.8 cells 36 hr], p=1.0]). To quantify the PCNA-positive cells for the 48-hr. time point, the mechanism changed from counting individual cells to counting cell clusters because at 48 hr., the progenitors had asymmetrically divided to form clusters of 2-4 cells. The clusters were significantly different between the DA and NDA groups, with 16.1±8.8 clusters and 25±18.1 clusters, respectively (Figure 4C, p=0.00509). In conclusion, dark adaptation is not needed during LIRD to produce robust damage and MG proliferation response, shortening protocols across the country, and eliminating the need for dark boxes to be placed on housing systems.

**Figure 4:**
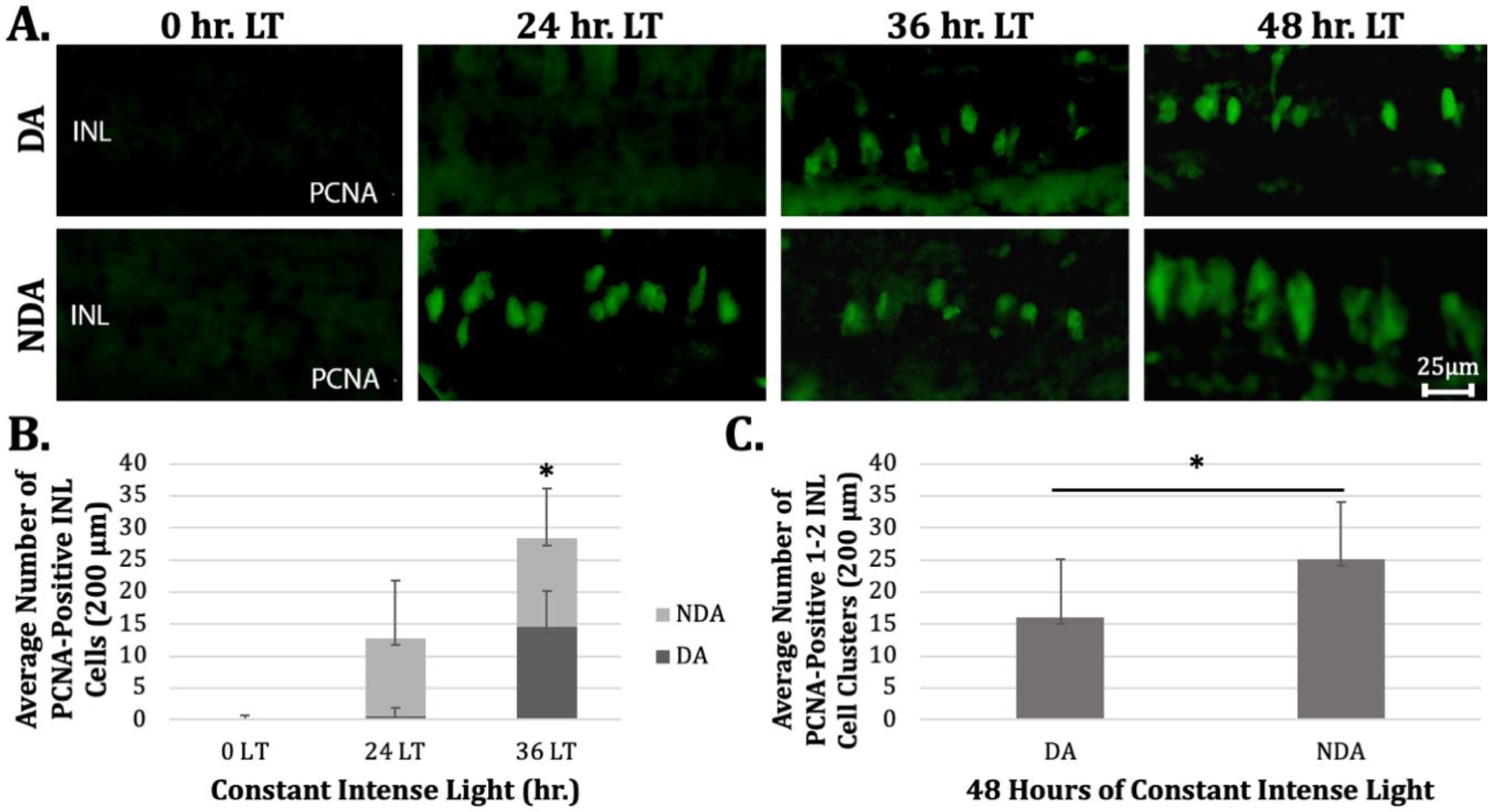
Light damage significantly increased INL PCNA+ cells in both dark-adapted and non-dark-adapted retinas. A) DA and NDA retinas were light damaged, sectioned, and immunohistochemically labeled for PCNA (green) to label proliferating Müller glia cells and progenitors-the hallmark of regeneration. B) For 0 hr., 24 hr., and 36 hr. light treatments, there were single cells that significantly increased in number as light damage occurred. Interestingly, there was a significant increase in NDA vs DA at 24 hr., with p=0.0229 indicating the paradigm worked. C) At 48 hr. light treatment, 2-4 cell clusters formed as the Müller glia divided. The number of PCNA-positive cells in NDA was significantly increased over DA, with p=0.00509. n≥10, p≤.05, ANOVA, Tukey post hoc.

## Discussion

This study presents an altered protocol design for LIRD, showing that the traditional 10–14-day dark adaptation period prior to LT is not necessary to drive a proliferation response in the adult zebrafish retina (Figure 3 A-C). It was previously thought that the dark adaptation period was needed to prime the retina to receive more light after being transferred to LT, therefore, causing more damage to the retina and generating a larger proliferative response. When the retina undergoes injury, a signal cascade is initiated through transcription factors and genetic expression that encourages Muller glia to re-enter the cell cycle and begin dividing to regenerate the retinal cells lost in the lesion (Figure 1C). As seen here, forgoing the dark adaptation period produces the same, and in some cases better, results than the zebrafish that were dark adapted.

To drive a proliferation response, damage must first occur in the retina. To verify a damage response, retinas were stained with DAPI, a nuclear dye, to visualize the ONL thickness (indication of integrity) and number of rod photoreceptor nuclei present. As damage continues, the retinal layers collapse in on each other. Figure 2A shows the obvious morphological disparities as the LT continues, particularly that the ONL, INL, and ganglion cell layer all collapse inward toward the midline as the structural integrity of the retina is lost and cells begin to die. In both DA and NDA, there is a significant decrease in ONL thickness and number of rod nuclei from 0 to 24 hr. (Figure 2B and 2C), indicating that LT caused damage in both models. There is no significant difference between the two groups, however, at any of the LT time points, suggesting dark adaptation is not needed to prime the retina to receive more light and produce a greater amount of damage, as previously thought. The only notable difference between the two groups is the shape of the NDA rod nuclei. Starting at 24 hr. LT and continuing through 48 hr., NDA rod nuclei take on a shape differing from that of DA rod nuclei. The NDA nuclei have a thickly rimmed shell with a light center, causing them to appear concaved (Figure 5). It is unknown if the shape of the nuclei affects the structure and/or function of the retina or what causes this shape to occur, and it is an area of interest for future study.

Across zebrafish research, there is not yet a standard expectation for cone cells in the way there is for rod cells in response to LIRD (D’Orazi et al., 2020; Khan et al., 2020; Vihtelic & Hyde, 2000). However, it is expected that the cones experience an overall decline in morphology. ZPR-1 was used to visualize red/green double cones. As seen in Figure 2A, the cones become less aligned, more dispersed, and they begin to fray at the ends with the disappearance of the end feet as LT increases. There was also a decrease in cones in both DA and NDA groups. The most severe decrease occurred from 0 to 24 hr., with DA decreasing from 45 to 33 cells and NDA decreasing from 39 to 18 cells. There was no significant decrease or increase throughout the rest of the time points in either group. Yet, from 0 to 24 hr., there is a significant difference between the two groups, with the NDA having a more dramatic and significant decrease in the number of cone cells compared to DA controls. The mechanism behind this difference is not yet understood. It is of note that, although not significant, there is a slight increase of 4 cells in NDA cone cells from 24 to 48 hr.

It is known that a threshold of damage must be achieved to induce a proliferative regeneration response (Vihtelic & Hyde, 2000). If dark adaptation is not required to structurally damage the retina, but there is no regeneration occurring, then changing the protocol is futile. As transcription factors and genetic expression change following lesion, Muller glia re-enter the cell cycle and begin to divide as multipotent progenitor cells that produce PCNA, which can then be used to visualize the progenitor cells as they divide and migrate in the INL. Quantifying the level of PCNA-positive cells in the INL indicates how responsive the retina was to the damage and its effort to repair. Insofar, the data has been similar between DA and NDA zebrafish, however, the INL PCNA begins to show discrepancies. The NDA retinas have an earlier and more robust proliferation response, displaying a significant increase from 0 to 12 cells during 0 to 24 hr. The DA has no change in INL PCNA-positive cells, remaining at 0 cells from 0 to 24 hr. However, from 24 to 36 hr., the DA cells significantly increased from 0 to 15 cells while the NDA showed a 1-cell increase over the same time points. To quantify the 48 hr. LT, cell clusters were counted instead of individual cells. As the progenitor cells asymmetrically divide and migrate, they form clusters of 2-4 cells that become difficult to differentiate as individuals. Due to this, the 48-hr. data is presented separately from the rest, in Figure 4C. There is a significant difference between DA and NDA clusters, with NDA having 25 clusters and DA having only 16 clusters. It is possible that the later and lesser proliferative response in DA retinas has to do with the dark adaptation itself. In this new model, there is no difference in the rod photoreceptor response, but the cones are damaged in a more robust fashion, which also correlates with a quicker and more robust Muller glia proliferation response. As the progenitor cells divide and migrate, they mature into precursors and travel to the ONL where they will further differentiate into photoreceptors. Presented in Figure 6, at 0 hr. there is a significant difference between DA and NDA, which is of note because these fish *should* be the experimental control group. However, DA retinas begin at 0 hr. with 7 ONL PCNA-positive precursor cells, while NDA has 0 as expected. Then, as LT continues, the NDA cells increase significantly from 0 to 6 cells during 0 to 36 hr. time points. It would be expected that there are far more ONL PCNA-positive cells at later time points as they migrate, so starting with such a high number of cells in the DA group suggests something is occurring due to the dark adaptation process. Oddly, at the 48-hr. time point, there is no significant difference between the DA and NDA group, each having 17 and 11 cell clusters, respectively. So, even though the proliferative response might be varying in ONL PCNA-positive cell numbers from 0 to 36 hr., it ends with similar responses that are not significantly different from one another, indicating that both models have sufficient regeneration in response to LIRD. More research is needed to elucidate the mechanism behind the proliferation response and to solve the puzzle the ONL PCNA-positive cell data presents.

## Acknowledgements

We would like to thank the Natural Science Fellows Program, the Biology Department, and the Neuroscience Program at High Point University for funding this work.

